# Investigating the quaternary structure of a homomultimeric catechol 1,2-dioxygenase: An integrative structural biology study

**DOI:** 10.1101/2024.12.05.627049

**Authors:** Arisbeth Guadalupe Almeida-Juarez, Shirish Chodankar, Liliana Pardo-López, Guadalupe Zavala-Padilla, Enrique Rudiño-Piñera

**Affiliations:** Laboratorio de Bioquímica Estructural,Departamento de Medicina Molecular y Bioprocesos, Instituto de Biotecnología, Universidad Nacional Autónoma de México, Cuernavaca, Morelos, México; National Synchrotron Light Source II, Brookhaven National Laboratory, Upton, New York, United States of America; Laboratorio de Biotecnología Marina, Departamento de Microbiología Molecular, Instituto de Biotecnología, Universidad Nacional Autónoma de México, Cuernavaca, Morelos, México; Unidad de Microscopía Electrónica, Instituto de Biotecnología, Universidad Nacional Autónoma de México, Cuernavaca, Morelos, México

**Author notes:** Corresponding author (ERP) and (AA).

## Abstract

The structural analysis of catechol 1,2 dioxygenase from *Stutzerimonas frequens* GOM2, SfC12DO, was conducted using various structural techniques. SEC-SAXS experiments revealed that SfC12DO, after lyophilization and reconstitution processes, can form multiple enzymatically active oligomers, including dimers, tetramers, and octamers. These findings differ from previous studies, which reported active dimers in homologous counterparts with available crystallographic structures, or trimers observed exclusively in solution for SfsC12DO and its homologous isoA C12DO from *Acinetobacter radioresistens* under low ionic strength conditions. In some cases, tetramers were also reported, such as for the *Rodococcus erythropolis* C12DO. The combined results of Small-Angle X-ray Scattering, Dynamic Light Scattering, and Transmission Electron Microscopy experiments provided additional insights into these active oligomers’ shape and molecular organization in an aqueous solution. These results highlight the oligomeric structural plasticity of SfC12DO, proving that it can exist in different oligomeric forms depending on the physicochemical characteristics of the solutions in which the experiments were performed. Remarkably, regardless of its oligomeric state, SfC12DO maintains its enzymatic activity even after prior lyophilization. All these characteristics make SfC12DO a very promising candidate for extensive bioremediation applications in polluted soils or waters.

## Introduction

Catechol 1,2-dioxygenases, C12DOs, constitute a class of non-heme iron-containing enzymes (Fe^+3^) in the family of intradiol dioxygenases (1). They play a crucial role in the □-ketoadipate pathway, specifically in the cleavage of the aromatic ring of catechol, by introducing two oxygen atoms between carbons one and two of the aromatic ring (2). This reaction leads to the formation of *cis-cis* muconate (*ccMA*), which subsequently enters the tricarboxylic acid cycle. Moreover, *ccMA* is a precursor of the adipic acid commonly used in the industry for the benzene-free synthesis of nylon 6,6 and other polymers (1,3,4).

C12DOs have been identified in many organisms, spanning bacteria, fungi, and even higher plants (1,5–11). Research on C12DOs has mainly focused on bacteria strains isolated from those substrates particularly affected by industrial residues, oil spills, and other contaminants (7). Some bacteria capable of surviving and growing in these highly polluted environments have been isolated, for example, *Gordonia alkanivorans*, obtained from the oil-contaminated sludge of a local gas station in Taiwan (12); *Paracoccus* sp. MKU1 isolated from the textile industrial effluent near Tirupur, India (13,14), and *Stutzerimonas frequens* GOM2 (*Pseudomonas stutzeri* GOM2) obtained from the oil-polluted substrate in the depth of southwestern Gulf of Mexico (15).

According to the crystallographic structures available in the PDB, native C12DOs are dimeric, as seen in PDB entries 2AZQ, 1DLM, 2XSR, 5UMH, 5TD3, 5VXT, and 3HGI. These dimers present a common hydrophobic zipper at the dimeric interface, with a pair of phospholipids embedded within the cavities of this interface (16). However, despite the dimeric homogeneity observed in crystallographic structures, some C12DOs present different oligomeric states in solution. For example, certain C12DOs exist as monomers, such as C12DO from *P. aeruginosa* TKU002 (M_W_ 22 kDa) (17), or trimers, such as the C12DO from *Trichosporon cutaneum* WY 2-2 (M_W_ 105 kDa)(18), and the C12DO from *Paracoccus* sp. MKU1-A ( M_W_ 121.4 kDa) (14).

In some cases, C12DOs enzymes also show a flexible quaternary structure. For example, C12DO from *Rhodococcus erythropolis* AN-13 is mainly described as monomeric, but in the absence of salt in the buffer, oligomerizes into stable tetramers (M_w_ 150 kDa) (19). The Isoform A of C12DO from *Acinetobacter radioresistens,* can exist in two active forms: as a trimer (M_W_ 112.4 kDa) and as a dimer (M_W_ 77.6 kDa), depending on the ionic strength conditions (20). For catechol 1,2 dioxygenase from *S.frequens* GOM2, SfC12DO, previously named PsC12DO, a transition from trimers into dimers was described when the buffer’s salt concentration mimics seawater (15). This intriguing behavior of oligomer transitions in some C12DOs implies that the biochemical environment, particularly the ionic strength, seems to be the critical factor driving those changes (15,20). Furthermore, these structural alterations may also be associated with variations in its enzymatic activity and the protein aggregation- dissociation behavior (15,19–22).

In this study, we utilized a combination of analytical techniques, including Small-Angle X-ray Scattering (SAXS), Size Exclusion Chromatography (SEC), Dynamic Light Scattering (DLS), and Transmission Electron Microscopy (TEM) to gain fundamental insights into the quaternary structure behavior of SfC12DO in solution. By understanding how structural changes influence enzymatic activity, we can better evaluate the potential application of these enzymes in bioremediation processes, including those involving seawater affected by oil spills. This work represents the first application of SAXS to SfC12DO, revealing new low-resolution structural characteristics of this catechol 1,2-dioxygenase, which exhibits several active oligomeric states in solution.

## Materials and Methods

### Protein Expression and Purification

To express SfC12DO we employed *E. coli* BL21 (DE3) cells harboring the pETCDOPs vector, which contains the CatA gene responsible for encoding catechol 1,2- dioxygenase derived from *S. frequens* GOM2. The details of the vector construction process are described in (15).

A pre-culture of *E. coli* BL21 (DE3) cells transformed with pETCDOPs was inoculated into 1L of LB medium containing 100 μg/mL ampicillin and grown at 37 °C, 200 rpm. Once the cell culture reached an OD_600_ of 0.6 - 0.7, protein expression was induced by adding IPTG at a final concentration of 1 mM and further incubated for 5 h. After this incubation, the cell culture was harvested by centrifugation at 4,600 *g* at 4 °C. The cells were disrupted by sonication at 37 % amplitude and 40 s ON / 40 s OFF cycle for 30 min (Sonics Vibra-cell Ultrasonic Processor), using *lysis Buffer* (S1 Table) throughout the sonication and purification processes. Affinity chromatography was conducted following three gradient steps of Imidazole (5 mM, 20 mM, and 500 mM). The SfC12DO purity was confirmed by SDS-PAGE (Sodium Dodecyl Sulfate - Polyacrylamide Gel Electrophoresis).

### Sample preparation

The purified SfC12DO was concentrated and filtered using Amicon® Ultra-15 centrifugal filters with a molecular weight cutoff of 30 kDa (Merk, Germany), undergoing three changes of 10 ml of MilliQ water at 4 °C.

Enzyme aliquots containing 5 and 10 mg were frozen at -80 °C and lyophilized for 12 hours using a LABCONCO Freezone® Legacy system. The lyophilized samples were then stored at -20 °C. Before analysis, the lyophilized SfsC12DO samples were reconstituted by adding Buffer B (S1 Table) to achieve the desired concentration for each experiment performed in this work.

### Circular Dichroism spectra (CD)

CD spectra of SfC12DO were acquired using a Jasco J-715 CD Spectrometer (JASCO Analytical Instruments) to assess the sample’s secondary structure composition. The protein concentration was adjusted to 0.3 mg/ml, with a sample volume of 200 μl. Two sample types-lyophilized-reconstituted and non-lyophilized SfC12DO were resuspended in degassed and filtered Buffer C (S1 Table). Measurements were performed in triplicate using a 1 mm pathlength cell over a wavelength range between 190 and 260 nm at 25 °C, controlled by a Peltier temperature cell holder (PTC-4235; JASCO). The ellipticity was reported as mean residue ellipticity ([θ]mre, in deg cm² dmol⁻¹), and the data were analyzed using the BestSel web server (http://bestsel.elte.hu).

### Enzyme activity assay

The specific activity of SfC12DO was determined spectrophotometrically using a Cary 60 UV-vis spectrophotometer (Agilent Technologies). The enzymatic reaction was monitored by measuring the formation of product (*ccMA*) at 260 nm (ε = 16.8 mM) (23) at 40 °C for 1 min. One unit (U) of the enzyme was defined as the amount of the enzyme required to catalyze the formation of 1 µmol of ccMA per min.

### Size Exclusion Chromatography (SEC)

For SEC analysis, an aliquot of SfC12DO of 500 μl volume at 10 mg/mL was centrifuged at 12,800 *g* for 5 min to eliminate aggregates. Following centrifugation, the sample was subjected to SEC using a Sephacryl 200 HiPrep 16/60 column on an ÄKTA Prime - FPLC System. The column was pre-equilibrated with Buffer B (S1 Table). The purification process was conducted at a 1.0 mL/min flow rate, and the absorbance at 280 nm was monitored. The molecular mass was confirmed by comparing the retention time to the calibration curve standards. Lysozyme (14 kDa), Horseradish (44 kDa), PADA-I* (52 kDa), BSA (66 kDa), and ɣ-globulins (150 kDa).

*PADA-I was kindly provided by Prof. Marcela Ayalás Group, IBt, UNAM. All other proteins used for column calibration were sourced from Sigma-Aldrich.

### Denaturing and Non-denaturing PAGE

*SDS-PAGE* was carried out by the denaturing Tris-Gly-SDS buffer system (Laemmli, 1970). A Precision Plus Protein™ prestained marker (Bio-Rad) was used as the standard (in the range from 10- to 250 kDa). Electrophoresis was performed at 120 V at room temperature, and gels were stained with Coomassie blue.

*SEMI-NATIVE-PAGE* was carried out using the Tris-Gly-SDS buffer system (Laemmli, 1970) without adding detergent or denaturing agents. BSA was used as a protein size standard. Electrophoresis was performed at 100 V at 18 °C, and gels were stained with Coomassie blue.

### Dynamic Light Scattering analysis (DLS)

DLS analysis was performed using the Zetasizer Nano Z system (Malvern Instruments) on a SfC12DO protein sample solution with a volume of 1 mL at 1 mg/mL. The measurements were carried out in a quartz cuvette, and the scattered light was collected at a fixed angle of 173 °. The experiments were performed in triplicate to ensure accuracy and reliability. The temperature during the analysis was maintained at 40 °C.

### Transmission Electron Microscopy (TEM)

SfC12DO sample solutions with concentrations in a range between 0.3 and 0.5 mg/mL were negatively stained using the methodology described in (24). Samples were applied onto EMS carbon copper grids and washed briefly in sterile water for 5 seconds. Uranyl acetate (1 % w/v) (Electron Microscopy Science) was used on the grid for 30 s, and the excess stain was blotted directly. Subsequently, the uranyl acetate application was left to dry for 5 min before being mounted in the sample holder. TEM was performed at 80 kV using a ZEISS LIBRA 120 Transmission Electron Microscope. Images were recorded with a GATAN CCD, and the data obtained was analyzed with the Digital Micrograph Software (Gatan, Pleasanton, CA).

### Structural modeling of SfC12DO

To obtain tridimensional models of SfC12DO based on its amino acid sequence, we used AlphaFold2 (25). Multimer analysis using the AlphaFold2-multimer was applied to model ten putative SfC12DO oligomers (monomer, dimer, and trimer) that were predicted using ColabFold v1.5.5 (https://colab.research.google.com/github/sokrypton/ColabFold/blob/main/AlphaFold 2.ipynb) (26) and tetramer to decamer models were generated through the COSMIC^2^ platform (https://cosmic2.sdsc.edu) (27). For each modeling run, five structures were generated and selected based on the top predicted local-distance difference test values (plDDT).

### Small-Angle X-ray Scattering (SAXS)

SAXS experiments were performed at the LIX-beamline (16-ID) of the National Synchrotron Light Source II (NSLS-II, Upton, NY). The SfC12DO sample was previously resuspended in Buffer B (S1 Table) and centrifuged. SAXS data analysis was analyzed using two different methods. For *static mode,* the buffer scattering was determined by subtracting the scattering of the empty cuvette from that of the degassed Buffer B (S1 Table). The protein and buffer scattering measurements were repeated in three separate experimental sessions, and the results were averaged. The data were processed by the ATSAS version 3.0.5-2 package (28). The data quality is assessed using AUTORG, and SAXS profiles were further analyzed with OLIGOMERS to interpret multi-component mixtures (30).

For SEC-SAXS (size exclusion chromatography coupled to X-ray scattering data collector) analysis, the sample was applied into a pre-equilibrated Superdex 200 5/150 column (GE Healthcare) with degassed Buffer B (S1 Table) as the mobile phase. The eluent was directly coupled to X-ray scattering data collection.

Data collection utilized a wavelength of 0.819 Å and a scattering angle of 0.006 < *q* < 3.0 Å^−1^, with an exposure time of 2 s. Scattering data was collected using Pilatus 3X 1M and 900 K detectors positioned at 3.7 m and 300 m sample-detector distances, respectively.

### SAXS data processing

Initial data processing was performed using LiXTools (https://github.com/NSLS-II-LIX/lixtools) and py4xs (https://github.com/NSLS-II-LIX/py4xs). The determinations of I (0), molecular weight (MW) (Bayesian inference method), maximum dimension particle (Dmax), and radius of gyration (Rg) were calculated using PRIMUS in the ATSAS version 3.0.5-2 package (28). Rg was calculated from the slope of the Guinier plot, described as ln(I) vs. *q*², where q = 4 π sin (θ) / λ is the scattering vector (2θ is the scattering angle and λ is the wavelength).

For *ab initio* modeling, WAXSiS (31) (https://waxsis.uni-saarland.de) was used to fit the SAXS data, considering the explicit solvent modeling of protein (31,32). The SfC12DO models were initially generated using AlphaFold2 and subsequently refined with SREFLEX (33). Following these steps, for single component mixture, the quality of experimental data related to the protein model and hydration shell fitting was assessed using CRYSOL (34) and FoXs (https://modbase.compbio.ucsf.edu/foxs/) (35,36). This methodology for SAXS processing data is detailed in **Fig 1**.

**Fig 1.**
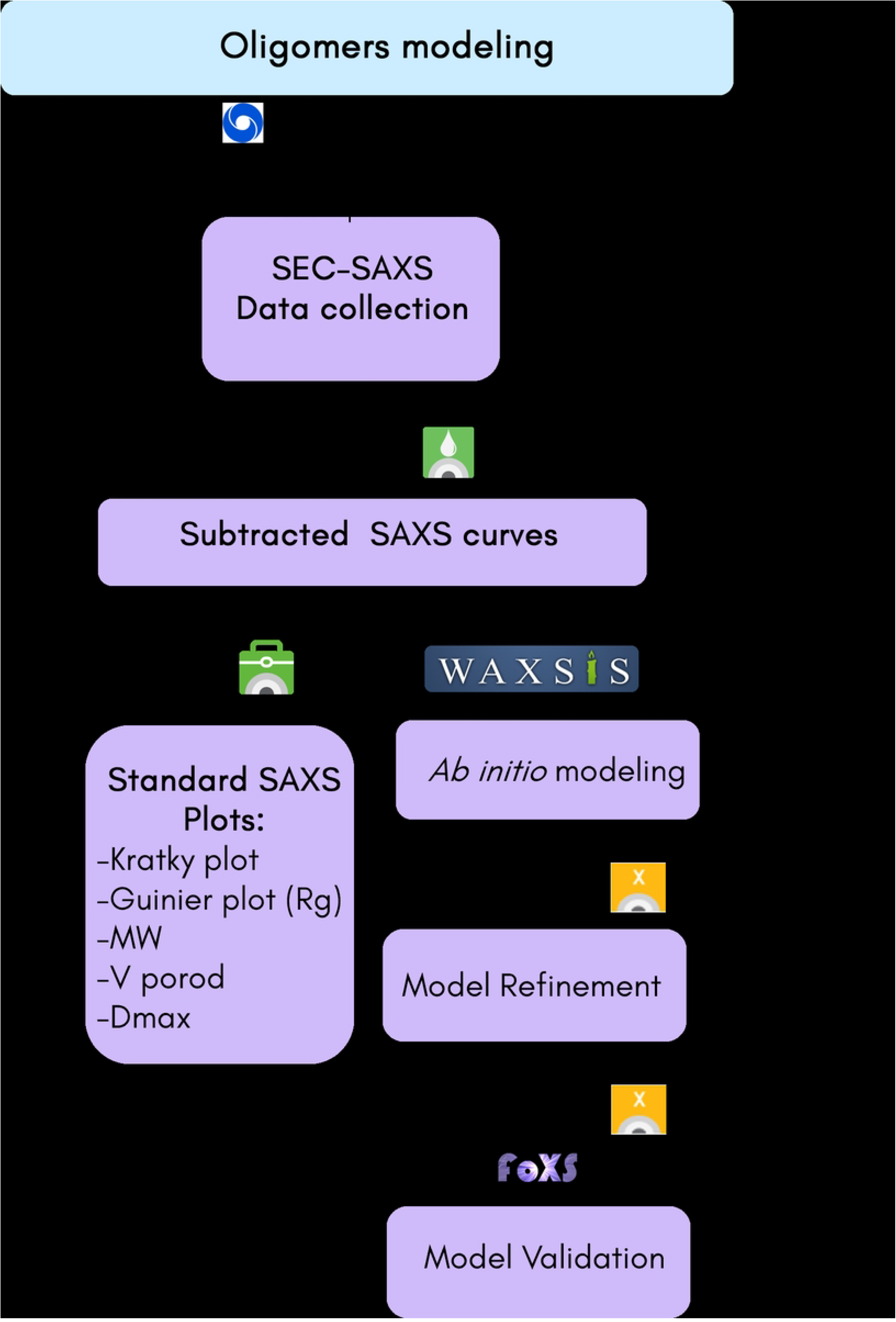
Schematic representation of SAXS data analysis process.

## Results and Discussion

### Secondary structure analysis

The CD analysis of SfC12DO’s showed slight changes in the content of secondary structure elements between non-lyophilized and lyophilized SfC12DO. The spectra reveal similar profiles with slight variations, particularly in the α-helix characteristic regions (∼208 nm and 222 nm) (Fig 2B). The distribution of α-helices and β-sheets closely resembled the typical C12DO secondary structure, showing significant similarity to Iso A (C12DO) from *A. radioresistens* S13 (2), which shares 53 % sequence identity with SfC12DO, and C12DO from *Achromobacter xylosoxidans* DN002, which has 50 % sequence identity with SfC12DO (37) (Fig 2C). Non-lyophilized SfC12DO contains 22.8 % α-helices and 27.5 % β-sheets, while lyophilization and reconstitution result in slight increases in α-helices (29.2 %) and decreases in β-sheets (26.2 %).

**Figure 2:**
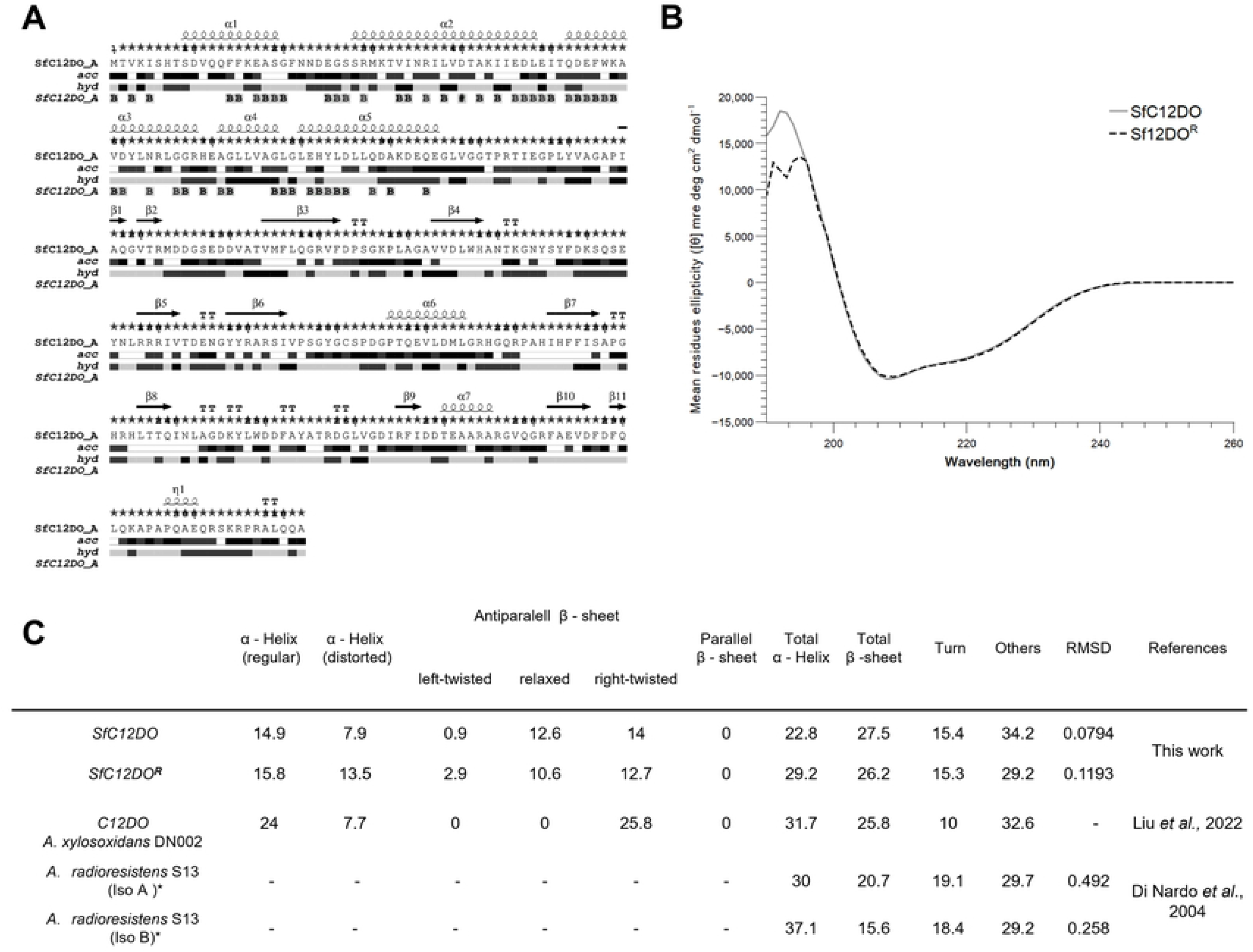
Secondary Structure of SfC12DO. **A)** The secondary structure of SfC12DO was predicted using ESpript 2.0. In this panel, *acc* represents the solvent accessibility index, and *hyd* stands for the hydrophobicity index **B)** CD spectra illustrate the content of the secondary structure in SfC12DO. The solid line represents non-lyophilized SfC12DO (gray), while the dashed line represents lyophilized and reconstituted SfC12DO^R^ (black). **C)** The table compares the content of secondary structure elements (α-helices, turns, coils, and β-sheets) in SfC12DO and SfC12DO^R^.

### SEC analysis and semi-native PAGE

The semi-native PAGE analysis of the lyophilized SfC12DO revealed the presence of various oligomers in contrast to the non-lyophilized SfC12DO, where only a dimer of approximately 70 kDa was detected (**Fig 3B**). Previous studies have reported the existence of various oligomers in the lyophilized C12DOs from *P. putida* (38); however, a detailed characterization of these oligomers is still needed to understand their structural and functional implications. The observed conformational differences are likely attributable to artifacts caused by the lyophilization process (38), which may promote non-specific aggregation or stabilize transient oligomeric forms that do not necessarily reflect the solution-state behavior of SfC12DO. The presence of distinct putative oligomers regarding the dimeric state was also confirmed by SAXS (*static mode*) OLIGOMERS (S2 Fig).

**Fig 3.**
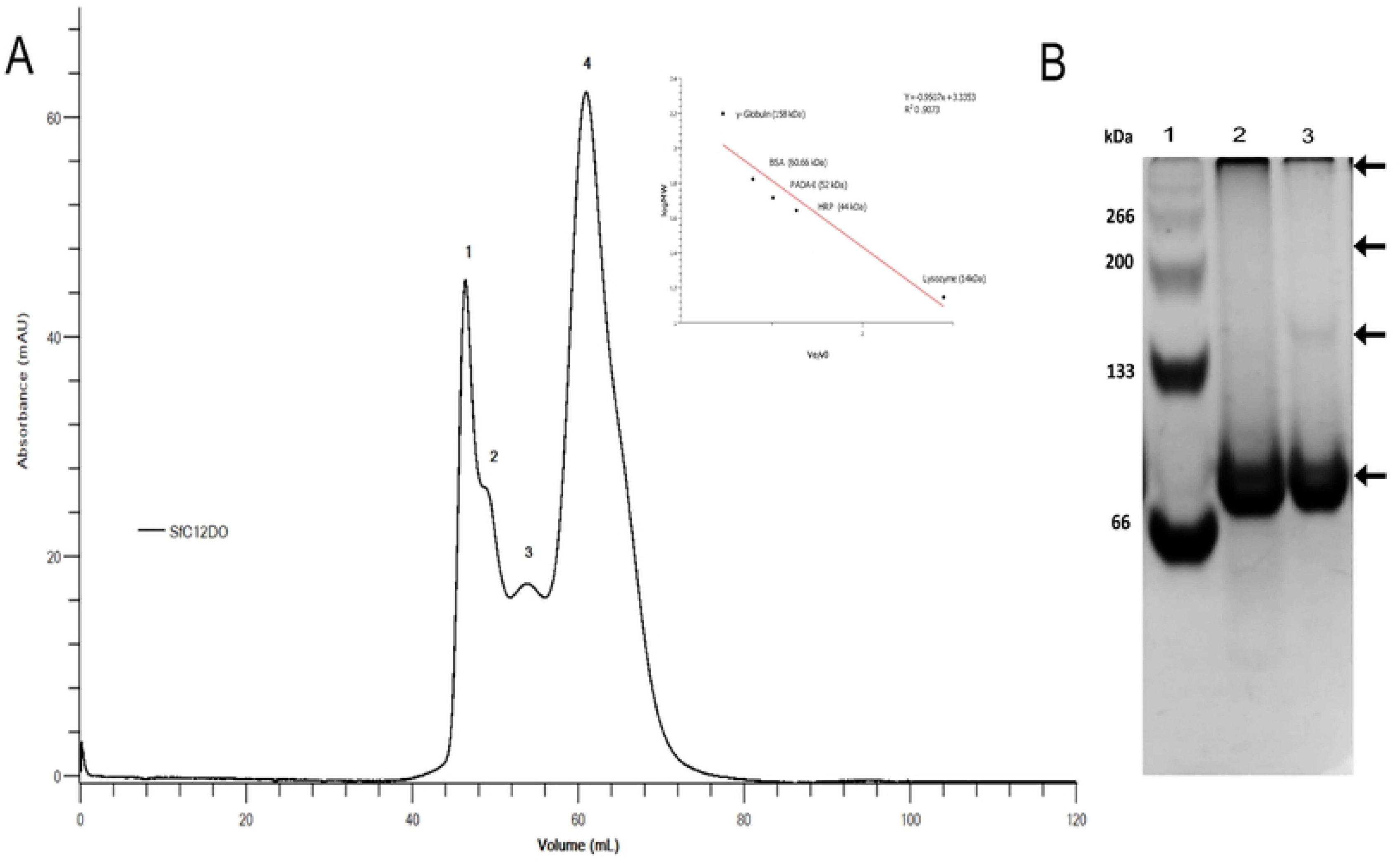
SEC elution profile of SfC12DO^R^ (lyophilized and reconstituted). A) SfC12DO was eluted from a Sephacryl 200 HiPrep 16/60 column. Four distinct peaks were observed: peak 1 (p1) with undetermined molecular weight (M_W_), peak 2 (p2) with undetermined M_W_, peak 3 (p3) with an estimated M_W_ of approximately 113 kDa, and peak 4 (p4) with a M_W_ of approximately 77 kDa. On the right, the calibration curve standard is shown. The black spots on the calibration curve represent protein standards. **B)** Semi-native-PAGE 8 % of SfC12DO conformations. Lane 1: BSA as a protein size marker. Lane 2: non-lyophilized SfC12DO. Lane 3: lyophilized and reconstituted SfC12DO protein.

To separate the different SfC12DO oligomers, a size-exclusion chromatography (SEC) analysis was performed (**Fig 3A**). The chromatogram exhibited four peaks at different retention volumes: p1 (46.5 ml), p2 (49 ml), p3 (54 ml), and p4 (61 ml). The calibration with protein standards enabled the estimation of molecular weights (M_w_) for peaks 3 and 4 (p3 and p4). However, the initial peak (p1) contained aggregates with molecular weights above exceeding the resolution limits of the SEC column employed, and the oligomers presented in p2 were bigger than the protein standard calibrators used in this experiment. The p3 content corresponds to an oligomer of approximately 113 kDa, and the p4 to a molecular weight of approximately 77 kDa, which resembled those M_w_ observed for the dimer in semi-native PAGE (**Fig 3**). A comparative analysis with protein standards unveiled the presence of three new distinct larger oligomers, corresponding to peaks p1, p2, and p3, the three surpassing the molecular weight of the trimeric (∼108 kDa) and the dimeric (∼80 kDa) SfC12DO reported in (15).

### SfC12DO activity assay

The enzymatic activity of four different peaks (p1, p2, p3, and p4) was evaluated using catechol as a substrate (**Fig 4**). Despite being lyophilized, peaks p3 (16.49 ± 1.2 U/mg) and p4 (15.42 ± 0.26) exhibited comparable activity to the non-lyophilized protein (15 ± 0.25 U/mg). This preservation of activity in C12DOs after lyophilization has also been observed in C12DO from *P. putida* DSM 437, where the lyophilized C12DO maintained its catalytic function in the presence of *n*-hexane (38). However, larger aggregates (p1 and p2) exhibited a significant reduction in specific activity, with a decrease of 81.6 % (2.75 ± 1.8 U/mg) and 63.4 % (5.49 ± 0.58 U/mg), respectively, compared to the non-lyophilized SfC12DO dimeric sample.

**Fig 4.**
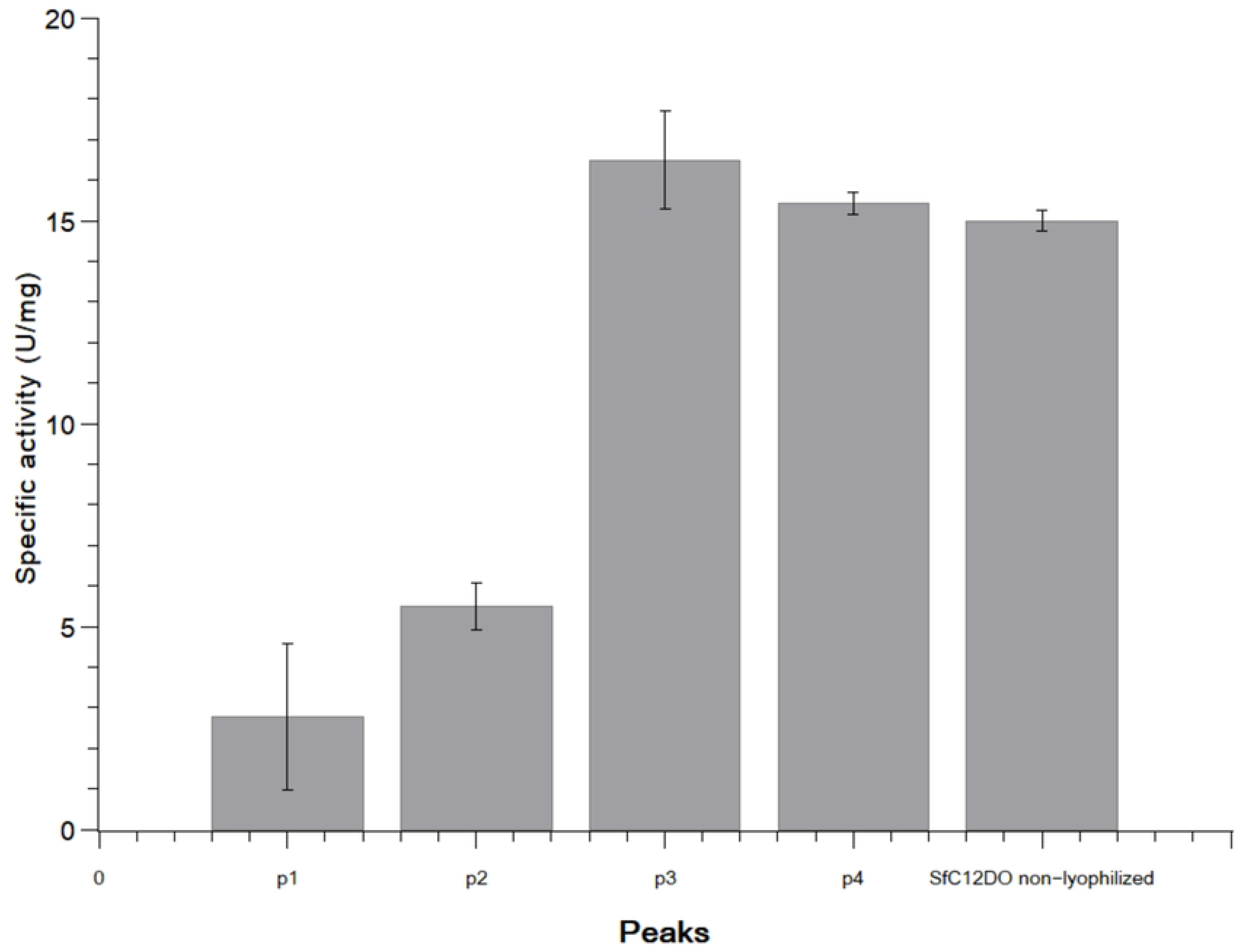
Specific enzymatic activity of SfC12DO (U/mg). The specific activity was monitored by measuring the formation of ccMA at 260 nm (ε = 16.8 M) at 40 °C for 1 min. One unit (U) of the enzyme was defined as the amount of the enzyme required to catalyze the formation of 1 µmol of ccMA per min. 1: p1 (2.75 ± 1.8 U/mg), 2: p2 (5.49 ± 0.58 U/mg), 3: p3 (16.49 ± 1.2 U/mg), 4: p4 (15.42 ± 0.26), 5: SfC12DO non-lyophilized (15 ±0.25 U/mg).

#### SEC - SAXS analysis

The four peaks obtained in SEC were subjected to SEC-SAXS data collection to further analyze the oligomerization of SfC12DO in solution. This revealed the structural organization and size of p1, p2, p3, and p4. **Fig 5** shows the intensity SAXS profiles for each of the four peaks of SfC12DO.

**Fig 5.**
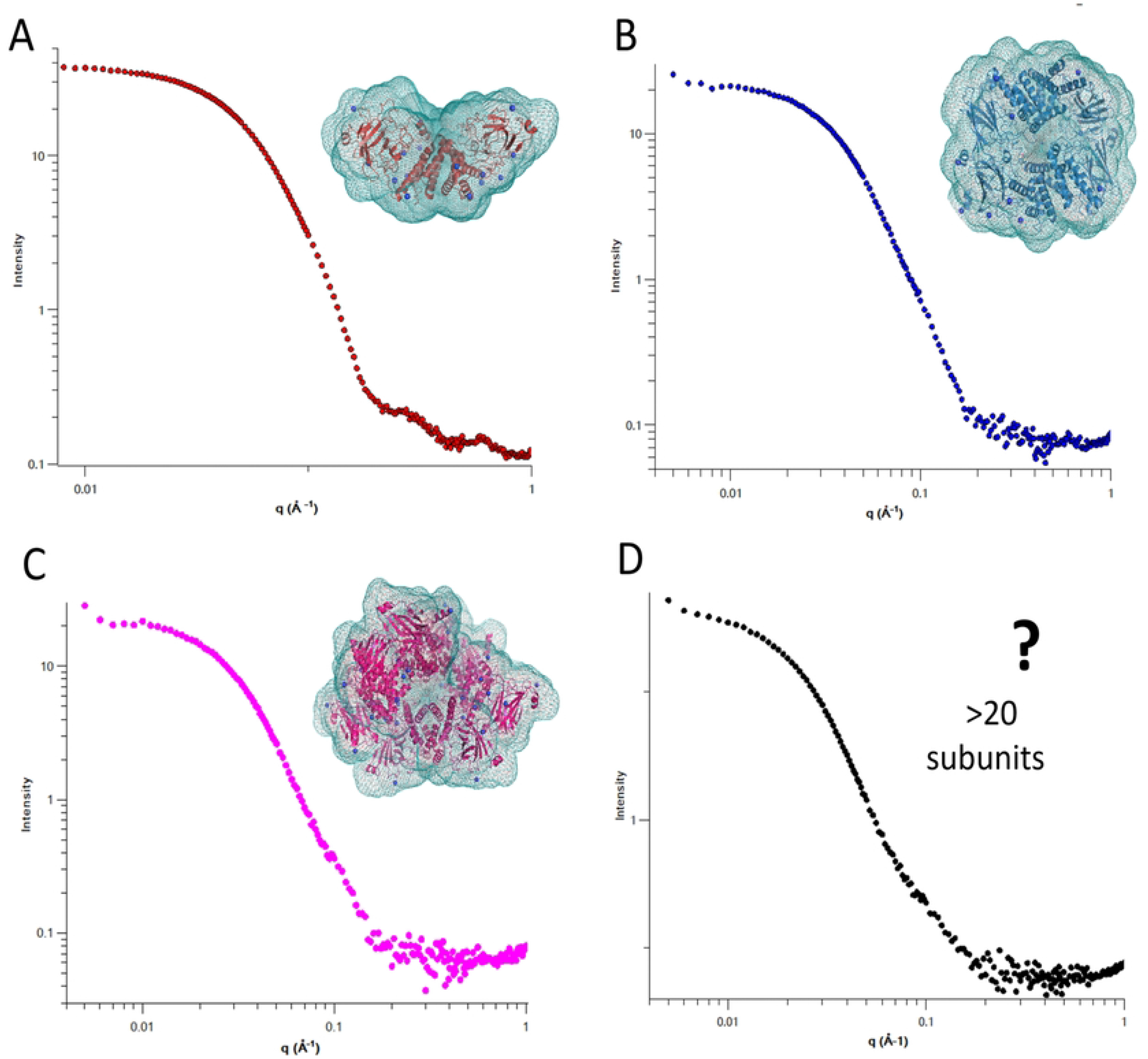
Integrative SAXS profiles of SfC12DO oligomers. Processed solution scattering intensity patterns of oligomers of p1, p2, p3, and p are shown. Each panel corresponds to different oligomeric states: A) Dimer, B) Tetramer, C) Octamer, D) Higher-order oligomer (> 20 subunits). The scattering profiles are plotted as intensity *versus* scattering vector (q), with the respective structural envelopes obtained through SAXS modeling.

The molecular weights of each oligomer were determined using Bayesian Inference (PRIMUS). The calculated molecular weights of p1 (833.40 kDa), p2 (318.45 kDa), p3 (157.05 kDa), and p4 (72.40 kDa) were utilized to classify the oligomeric states present in the SfC12DO sample. Given that the monomeric unit of SfC12DO has a molecular weight of 35.76 kDa, it can be deduced that p1 corresponds to aggregates with more than 20 subunits, p2 is an octamer, p3 is a tetramer, and p4 is a dimer. Differences between the apparent molecular weights of oligomers analyzed by SEC and the theoretical molecular weight of the SfC12DO monomer (35.76 kDa) may be due to structural modifications in SfC12DO’s secondary structure or, more likely, to flexibility in certain protein regions as indicated by the Kratky plots and CD spectra analyses. Table 1 contains all experimental SAXS parameters.

**Table 1.** SAXS data collection and analysis parameters.

Discrepancies were observed between the Rg values determined by ATSAS and WAXSiS analyses, possibly due to the hydration shell considered in the WAXSiS calculation. The Rg values for the four different oligomers were calculated: for ATSAS, p1 (83.39 Å), p2 (59.59 Å), p3 (45.16 Å), and p4 (32.10 Å), while for WAXSiS, the Rg values were p1 (not calculated because the lack of a three-dimensional model), p2 (48.79 Å), p3 (35.97 Å), and p4 (31.92 Å). Guinier plots for all oligomers correlated well with the Rg values obtained through the distance distribution function, P(r), analysis **(S3 Fig**). *P(r)* results indicated Rmax range values from 110 to 247 Å. The Kratky plot displayed a semi-bell-shaped peak, indicative of a partially intrinsically disordered structure in SfC12DO (S4 Fig).

The three-dimensional models of the three oligomers of SfC12DO, generated using WAXSiS based on SAXS data, reveal a predominantly globular shape. The experimental SAXS data indicate that the dimer adopts a more extended conformation than that observed in both, the crystallographic structures of homologs deposited in the PDB (Entries: 2AZQ, 1DLM, 2XSR, 5UMH, 5TD3, 5VXT, and 3HGI), and the models predicted by AlphaFold2. When scoring 10,000 conformers against SAXS datasets with CRYSOL, the best-fitting structure clusters displayed an elongated N- terminus for the dimeric SfC12DO SAXS three-dimensional model.

The obtained three-dimensional model for SfC12DO dimer indicates that the SAXS dimer model has a wider open angle between the N-terminal and other regions labeled in this work as region I (residues 1-28), region II (residues 197-222), region III (residues 268-282), and region IV (residues 293-312) in the solution state compared to its crystallographic counterparts (**Fig 6**). These differences were previously named "variability regions" in SfC12DO (15). The root-mean-square deviations (RMSD) analysis showing these analyses is presented in **Fig 2C**

**Fig 6.**
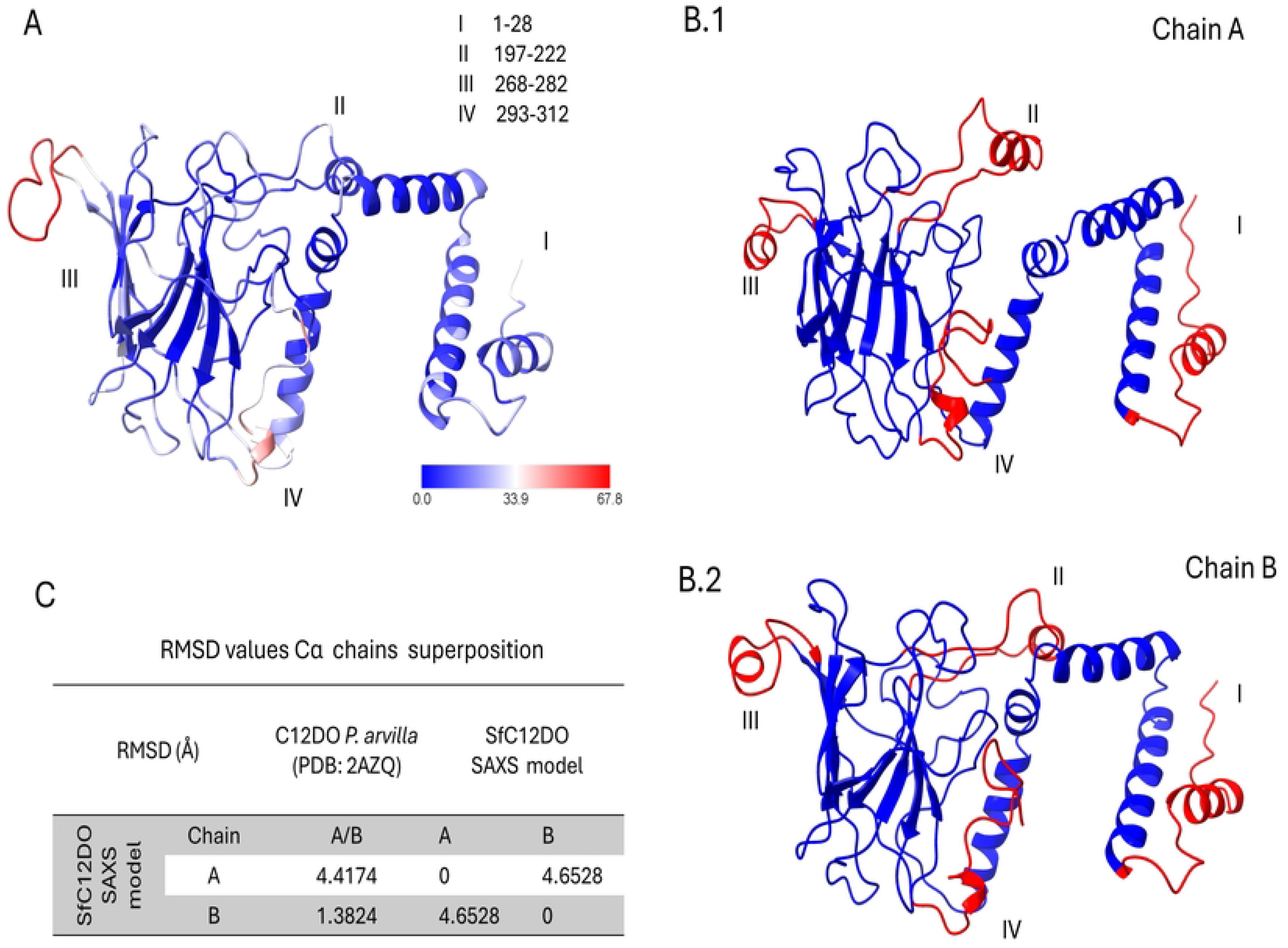
Comparative analysis of homologous C12DO with SfC12DO structure and flexible regions. A) The structure of the homologous C12DO (2AZQ) from *P. putida* shows that regions III and IV are flexible (colored in red). The structural coloring reflects the B values, with blue indicating lower flexibility (lower B values) and red representing higher flexibility (higher B values). Four key regions (I-IV) with varying flexibility are annotated: Region I (residues 1-28), region II (197-222), region III (268-282), and region IV (293-312). **B.1**) Chain A from the SAXS model exhibits regions of significant flexibility, highlighted in red, primarily regions I and III. **B.2)** Chain B follows a similar pattern but shows similarity to C12DO (2AZQ) in region II. D) Root Mean Square Deviation (RMSD) values show a comparison between *P. arvilla* C12DO (PDB: 2AZQ) and SfC12DO SAXS models for chains A and B. The results highlight structural differences between the models, with chain-specific deviations detailed

These differences could arise from the inherent flexibility of these regions in solution, allowing them to take on multiple conformations as observed in SAXS, but being restricted in their crystallized form. Such variations emphasize the importance of using both methods to fully comprehend the different shapes a C12DO can adopt (39). This may also explain how SfC12DO or certain C12DO oligomerizes, as they can take various structural arrangements without affecting their enzymatic activity.

The SAXS model of SfC12DO in tetramer and octamer form reveals previously undescribed conformations for C12DOs (**Fig 7**). These two SAXS conformations are likely possible and active in solution. In contrast, diffusion problems might prevent it from efficiently reaching all interaction target sites due to its size, negatively affecting the enzymatic activity in the octameric structure. Given the differences between X-ray crystallography and SAXS techniques in terms of structural detail and experimental conditions, the results, while distinct, indicate that the crystallographic dimer (like the dimer presented in the PDB entry 2AZQ) is present within the tetramer (**Fig 7A**). In the octamer, an arrangement of different SfC12DOs dimeric conformations is observed (**Fig 7B**).

**Fig 7.**
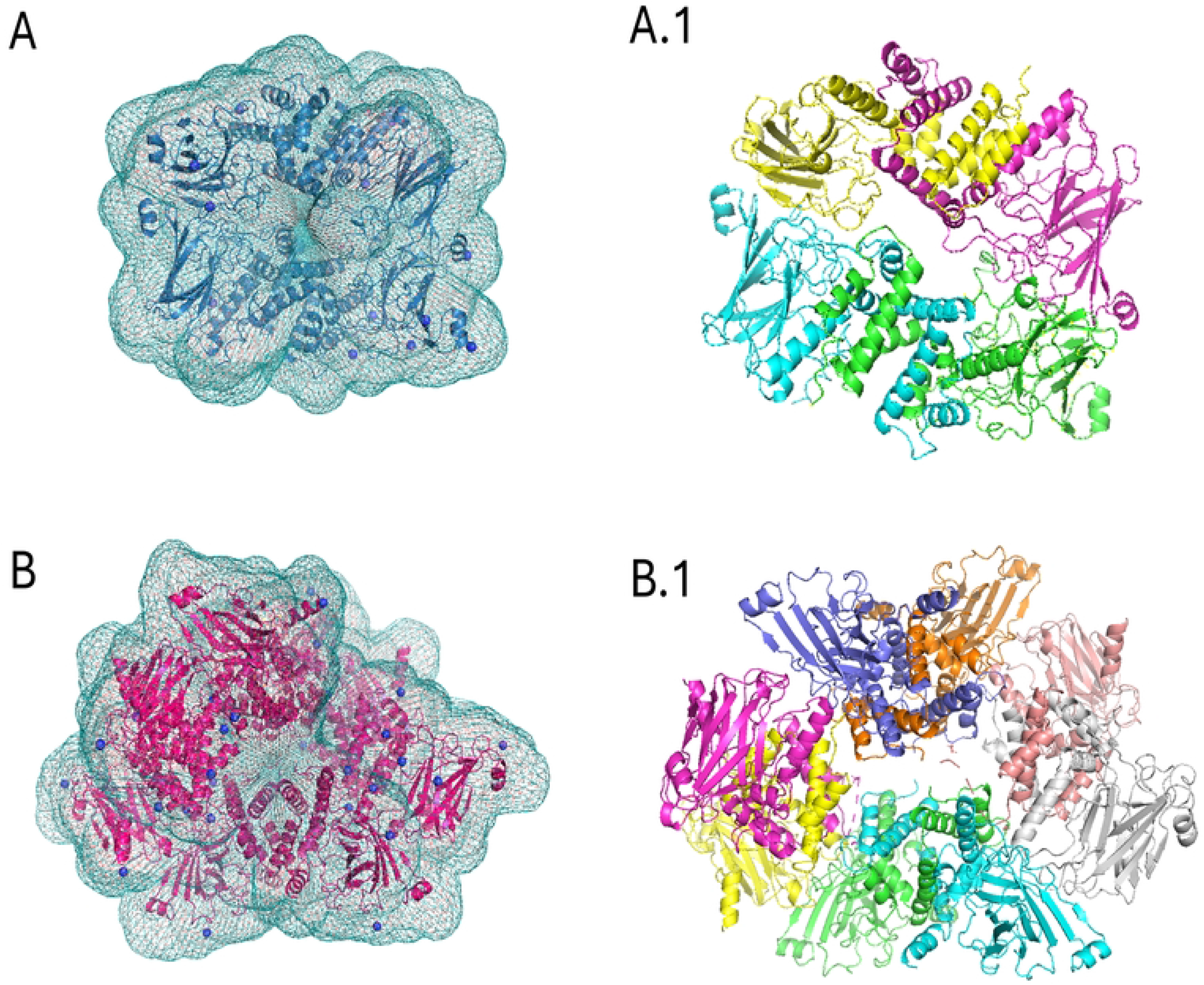
SAXS models for SfC12DO tetramer and octamer. A) SAXS envelope of the SfC12DO tetramer superimposed with its model. A.1) Cartoon representation of the tetramer, with each subunit shown in a different color B) SAXS envelope of the SfC12DO octamer superimposed with its model, respectively. B.1) Octamer structure with each subunit represented in a different color. Models were refined using SREFLEX AND DAMMIN.

### Characterization of oligomers by TEM and DLS

DLS initially evaluated the protein aggregation, and the calculated hydrodynamic diameters were between 7 and 20 nm. Subsequently, the determination of higher oligomers was confirmed by Transmission Electron Microscopy (TEM) (**Fig 8**). The resulting images using negatively stained contrast particles of different sizes correspond to the oligomeric species eluted in the SEC experiments (sizes from 75 kDa to 800 kDa). Due to the considerable variation in size, a detailed analysis using single-particle methods was not attempted. Nevertheless, a comparative study of TEM particles allowed for estimating their diameter. The TEM estimated particle diameter was consistent with the results obtained from SAXS.

**Fig 8.**
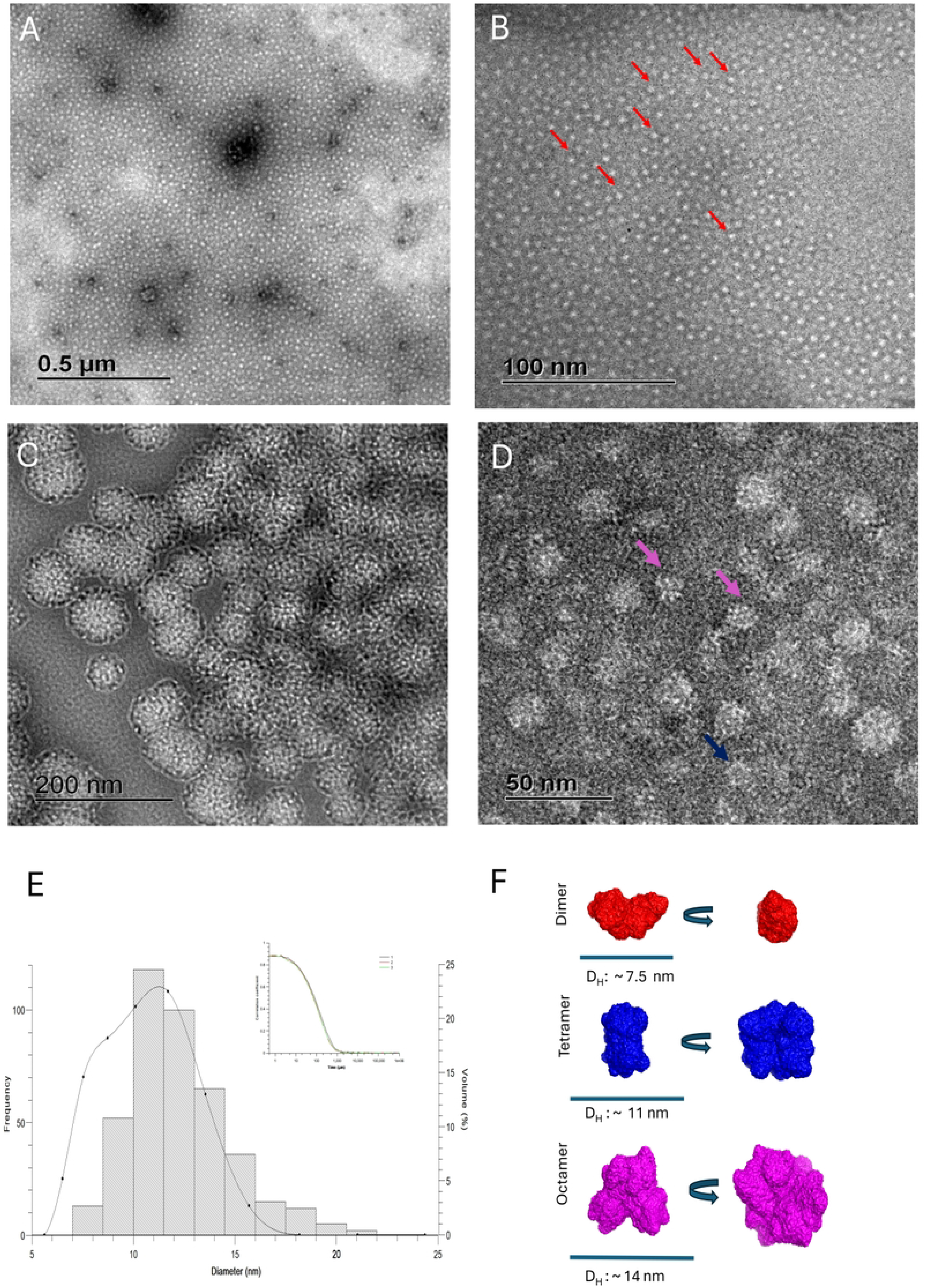
Negatively stained micrographs of oligomerized SfC12DO. (A-D) Transmission electron microscopy (TEM) images of SfC12DO oligomers at different magnifications, showing the structural organization of dimeric, tetrameric, octameric, and >20 subunits forms A). Overview of the sample with low magnification 12,000 X (scale bar: 0.5 0.5 μm). (B) The particles appear uniform at higher magnification, as seen at 80,000X (scale bar: 100 nm). (C) Aggregates of oligomers at intermediate magnification 31,500 X (scale bar: 200 nm). (D) Close-up of individual oligomers displaying different particle sizes and shapes corresponding to dimers (red), tetramers (blue), and octamers (pink) at 12,000 X (scale bar: 50 nm). All samples were prepared with 10 mM Tris HCl pH 8. The protein concentrations were 0.3 mg/mL, with an incubation of 5 min (E) Dynamic light scattering (DLS) analysis of SfC12DO oligomers, showing a histogram of particle size distribution with hydrodynamic diameters (D_H_) corresponding to a dimer (∼7.5 nm), a tetramer (∼11 nm), and an octamer (∼14 nm). The inset represents the autocorrelation function of the scattering intensity. (F) Model representation of SfC12DO oligomers based on SAXS. Dimers (red), tetramers (blue), and octamers (pink) are shown along with their respective hydrodynamic diameters (D_H_).

The SAXS-derived structural parameters mirror the SEC, native electrophoresis, TEM, and DLS results, consistent with different enzymatically active oligomeric forms of SfC12DO described in this work. These are highly unusual findings, given that other C12DOs have yet to be studied by SAXS and other techniques that evaluated their oligomeric behavior in solution at the level described in this work. Our results suggest that our SfC12DO preparations are composed of various functional oligomers that retain, at different levels, their functional activity in solution.

In this study, the dimeric form was the only SfC12DO oligomeric state consistently present across all the experimental approaches analyzed. This contrasts with the trimer form, proposed to appear only under certain optimal conditions (low ionic strength conditions) (14–16, 20, 40). Interestingly, the trimeric SfC12DO form did not appear after the lyophilization process nor in the SEC-SAXS results, suggesting that it might be a transient or a less stable form of SfC12DO or the experimental conditions used might disrupt the structural or specific conditions required for the trimer formation. These observations underscore the substantial impact of ionic strength on the formation and stability of different oligomeric states of SfC12DO, providing valuable insights into its structural behavior in diverse environments. More importantly, it demonstrates that oligomeric transitions in SfC12DO, and potentially in other C12DOs, are more common in solution, with several oligomers maintaining their enzymatic activity. This study highlights the crucial role of experimental conditions in determining the oligomeric states of proteins and provides a deeper understanding of SfC12DO’s structural dynamics and its potential implications in catalysis. Stressing the remarkable quaternary structure plasticity of C12DOs in specific environmental conditions and contributing to a more dynamic vision of the structural biology of proteins necessary for different processes, from basic to applied, to solve problems like bioremediation of soils and water. They highlight the remarkable quaternary structure plasticity of SfC12DOs in specific environmental conditions, which is crucial in understanding their structural dynamics and potential implications in catalysis.

## Acknowledgments

We want to express our gratitude to B.S. Maricela Olvera-Rodríguez, Biol. Rosa Roman-Miranda and Ph.D. Paloma C. Gil-Rodríguez for their technical assistance. We also thank Professor Gloria Saab-Rincón’s group for helping with Circular Dichroism (CD) measurements. Special thanks to Professor Marcela Ayalás Group at the Instituto de Biotecnología, UNAM, particularly MSc. Alina E. Torres-Aguirre, for kindly providing a sample of PADA-I used in the SEC-column calibration. Additionally, we acknowledge the Unidad de Microscopía Electrónica (UME) at the Instituto de Biotecnología, UNAM, for conducting the Transmission Electron Microscopy (TEM) experiments presented in this work.

